# Protection of high-frequency low-intensity pulsed electric fields and BDNF for SH-SY5Y cells against hydrogen peroxide-induced cell damage

**DOI:** 10.1101/2022.05.10.491265

**Authors:** Guan-Bo Lin, Wei-Ting Chen, Yu-Yi Kuo, You-Ming Chen, Hsu-Hsiang Liu, Chih-Yu Chao

## Abstract

Neurodegenerative diseases pose a significant global health threat. In particular, Alzheimer’s disease, the most common type causing dementia, remains an incurable disease. Alzheimer’s disease is thought to be associated with an imbalance of reactive oxygen species (ROS) in neurons, and scientists considered ROS modulation as a promising strategy for novel remedies. In the study, human neural cell line SH-SY5Y was used in probing the effect of combining non-invasive high-frequency low-intensity pulsed electric field (H-LIPEF) and brain-derived neurotrophic factor (BDNF) in protection against hydrogen peroxide (H_2_O_2_)-induced neuron damage. Our result finds that the combination approach has intensified the neuroprotective effect significantly, perhaps due to H-LIPEF and BDNF synergistically increasing the expression level of the phosphorylated epidermal growth factor receptor (p-EGFR), which induces the survival-related mitogen-activated protein kinases (MAPK) proteins. The study confirmed the activation of extracellular signal-regulated kinase (ERK) and the downstream pro-survival and antioxidant proteins as the mechanism underlying neuron protection. These findings highlighted the potential of H-LIPEF combined with BDNF in the treatment of neurodegenerative diseases. Furthermore, BDNF-mimetic drugs combining with non-invasive H-LIPEF to patients is a promising approach worthy of further research. This points to strategies for selecting drugs to cooperate with electric fields in treating neurodegenerative disorders.

## INTRODUCTION

In line with the aging of global population, neurodegenerative diseases (NDDs) have emerged as a mainstream medical issue, not only causing discomfort and even death for patients but also disrupting the life of their families. Among these brain diseases, Alzheimer’s disease is the most common type and a major cause of dementia in aged people. Alzheimer’s pathogenic mechanism is believed to be related to the aggregation of β-amyloid (Aβ), which leads to amyloid plaques, causing neuronal death via the generation of reactive oxygen species (ROS) [Chen and Zhong, 2014]. Another well-known hypothesis is phosphorylation of tau protein, which forms neurofibrillary tangles in nerve cell, damaging nerve cells and leading to progressive brain degeneration and deterioration of cognitive function [Spillantini and Goedert, 1998]. Although Alzheimer’s disease has been discovered for many years, there has yet to be an effective treatment, except a few therapies to relieve its symptoms. Hence, there has been a global scramble for the development of effective Alzheimer’s disease therapy. Neurotrophins are potential therapeutic agents for treating neurodegenerative disorder of the central nervous system. Among these neurotrophins, brain-derived neurotrophic factor (BDNF) is the most abundant neurotrophin in the human brain, which has a remarkable capability to repair brain damage. With proven effect in promoting neuronal survival, synaptic plasticity and neurogenesis, BDNF can resist brain damage caused by oxidative inflammation-related factors [Bathina and Das, 2015]. Some studies have found that BDNF produces neuroprotective effects against ROS-induced cell death, even with the potential for treating NDDs [Chen et al., 2017; Habtemariam, 2018; Palasz et al, 2020]. However, it should be noted that reduction in BDNF concentration can cause symptoms such as forgetfulness and poor learning ability, and many studies have blamed insufficient BDNF contents for depression and even Alzheimer’s disease [ Laske et al.,2006; Lee and Kim, 2009; Ng et al., 2019]. In addition, there has been evidence suggesting that Aβ can inhibit the proteolytic conversion of pro-BDNF to BDNF, decreasing BDNF level [Zheng et al., 2010]. Therefore, stimulation of BDNF secretion or use of exogenous BDNF is a promising approach in the treatment of Alzheimer’s disease. Many studies have proposed treatments capable of increasing the production of endogenous BDNF, thereby achieving nerves protection [ Hoppe et al., 2013; Wang and Holsinger, 2018; Uddin et al., 2020]. Meanwhile, the use of exogenous BDNF is also a direct and effective approach in dealing with the abnormal secretion of endogenous BDNF caused by the disease, as some studies have confirmed the significant neuroprotection effect of exogenous BDNF [Arancibia et al., 2008; Tapia-Arancibia et al., 2008; Xu et al., 2018]. However, a major problem is the short half-life of BDNF, which increases the difficulty of many therapeutic applications [Poduslo and Curran, 1996]. Therefore, how to improve BDNF efficiency will be an important issue.

In addition to traditional drug medication, physical therapy has gradually emerged as a mainstream approach in many diseases, such as electric current stimulation with proven value in treating muscle injury, wound, cancer, and neuronal diseases [Ainsworth et al., 2006; Perlmutter and Mink, 2006; Thakral et al., 2013; Wagstaff et al., 2016]. Many studies have pointed out that external electric stimulation can stimulate proteins to change their structure, affect their distribution, and even influence relevant biochemical pathways [Lamprea et al., 2002; Hekstra et al., 2016; Kawamura and Kano, 2019]. On the other hand, it is worth noting that epidermal growth factor receptor (EGFR) is a transmembrane protein deemed to be relevant to cell survival pathways. Especially, EGFR signalling abnormalities are believed to be associated with certain diseases, such as atherosclerosis, Parkinson’s disease, and Alzheimer’s disease [Dreux et al., 2006; Romano and Bucci, 2020]. Furthermore, it has been reported that the conventional electric current stimulation can affect cell migration via EGFR and enhance the expressions of downstream survival-related proteins, such as mitogen-activated protein kinases (MAPK) and phosphatidylinositol 3-kinase (PI3K) protein families in some medical fields [Chen et al., 2019].

To date, some issues remain unsolved for the application of electric current stimulation in many diseases. As the electric current stimulations are applied via contact or even invasive method, electric current would pass through tissues directly, causing the Joule effect [Novac et al., 2014]. It is known that invasive treatments are at higher risk and difficult in execution for some tissues, especially brain. Therefore, the development of non-invasive and non-contact electrotherapy is an inevitable trend. Recently, our team has developed a unique non-invasive and non-contact low-intensity pulsed electric field (LIPEF) stimulation for biomedical application. The pulsed electric field (PEF) can be considered as a composition of multiple sinusoidal subcomponents with different frequencies and intensities, which may interact with molecules, proteins, or organelles to induce specific bioeffect at the same time. Consequently, PEF stimulation can facilitate significantly the development of new therapies to meet different needs of different diseases, with great potential for applications in various medical fields. First, our team has demonstrated, for the first time, the capability of LIPEF to combine with epigallocatechin-3-gallate (EGCG) in inhibiting the pancreatic cancer cell line PANC-1 [Hsieh et al., 2017]. Second, low-dose curcumin, along with LIPEF, also has good anticancer effect, underscoring LIPEF’s capability to work with various chemical molecules [Lu et al., 2018]. Third, ultrasound can be added to the combination treatment, further boosting the anticancer effect [Hsieh et al., 2018]. Moreover, our team has found another effective anticancer triple therapy, combining LIPEF, chlorogenic acid (CGA), and thermal cycling-hyperthermia technique [Lu et al., 2020]. Besides, LIPEF can join hands with fucoidan to modulate rho-associated protein kinase (ROCK) and the downstream Akt protein, thereby protecting motor neurons NSC-34 against damage by oxidative stress [Hsieh et al., 2019]. On the other hand, in human neuronal SH-SY5Y cells under hydrogen peroxide (H_2_O_2_) stress, we found that high-frequency LIPEF (H-LIPEF) can modulate and activate the extracellular signal-regulated kinase (ERK) pathway, thereby protecting neuronal cells against oxidative stress [Chen et al., 2021]. In sum, in addition to safety, LIPEF (or H-LIPEF) alone or in combination with chemical molecules and/or other physical stimuli has significant potential for a wide range of applications.

In this paper, our team first combined the features of BDNF and H-LIPEF stimulation to study the effect of combination treatment on neuroprotection against the oxidative insult caused by H_2_O_2_ to SH-SY5Y human neural cells. The result showed that the combination of BDNF and H-LIPEF significantly inhibited the H_2_O_2_-induced cytotoxicity in SH-SY5Y cells. Examination of the underlying mechanism also indicated that the addition of BDNF is beneficial via regulation of ERK, a member of MAPK family proteins, a possible reason for the enhanced neuroprotective effect of combination BDNF and H-LIPEF treatment. Moreover, the combination treatment was found to significantly upregulate the expression of phosphorylated EGFR (p-EGFR) protein. These findings suggest that the combination of neurotrophic-related chemical molecules and H-LIPEF promises to be an effective therapy for neurodegenerative diseases.

## RESULTS

### In vitro-applied H-LIPEF

To avoid the harmful effects caused by conduction current and Joule effect, the culture dishes were placed in the non-invasive H-LIPEF device as described previously [Chen et al., 2021]. The H-LIPEF stimulation was performed in a non-contact manner with the apparatus consisting of two parallel conductive electrodes mounted on the insulating acrylic frame (Fig. 1A). Therefore, the H-LIPEF device can provide the electric field stimulation in a non-invasive manner, so that the cells were not in contact with the electrodes directly. The pulsed electric signal was from a function generator and amplified by a voltage amplifier that made pulsed electric signal tunable. The applied electric signal was characterized by a pulse train waveform (Fig. 1B) with various field strengths and frequencies for different applications. In this study, we used the electric field strength of 10 V/cm at 200 Hz, which was adopted in our previous study to be the optimal H-LIPEF condition for neuroprotection on SH-SY5Y cell line [Chen et al., 2021].

**Fig. 1.**
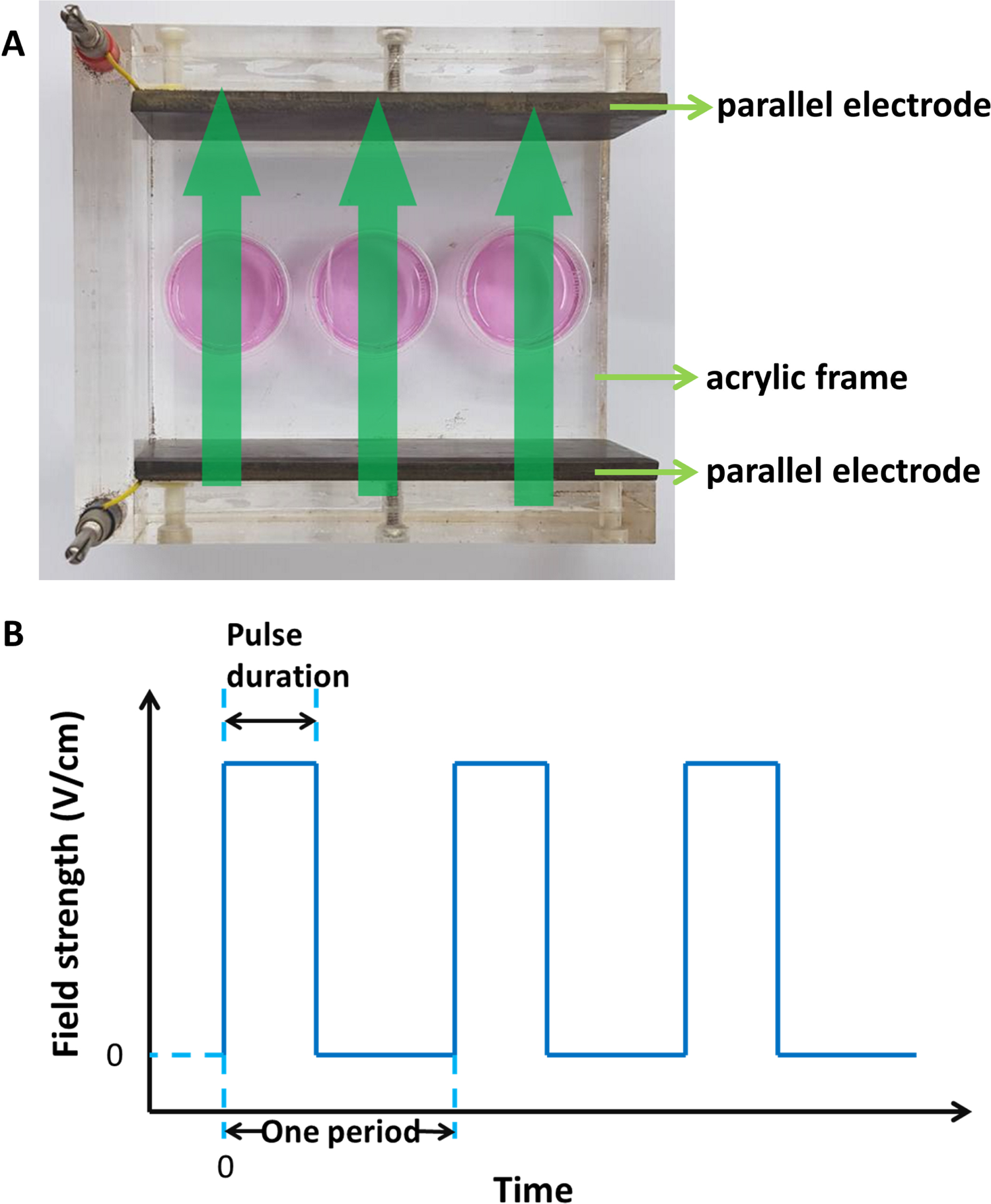
The experimental setup of H-LIPEF. (A) Image of the H-LIPEF stimulation device. The SH-SY5Y cells were placed in the H-LIPEF stimulation device constructed of two parallel electrodes. The green arrows shown here indicate the schematic representation of non-invasive and non-contact H-LIPEF exposure on cells. (B) Schematic representation of the pulse train waveform applied across the electrodes.

### Effect of the H-LIPEF and BDNF on H_2_O_2_-induced cytotoxicity in SH-SY5Y cells

Because oxidative stress plays a key role in the etiology of neurodegenerative diseases, reducing oxidative stress is often a top priority for neuroprotection. As shown in Fig. 2A, administration of H_2_O_2_ on SH-SY5Y cells for 24 h decreased the cell viability in a concentration-dependent manner. Treatment of SH-SY5Y cells with 500 μ for 24 h reduced cell viability to about 50% compared to the untreated control group. In order to understand whether the H-LIPEF combined with BDNF could attenuate the H_2_O_2_-induced neuron cell damage, we first applied various concentrations of BDNF (0 to 100 ng/ml) in SH-SY5Y cells and see their neuroprotection effect. As shown in Fig. 2B, we found a slight protective effect of these concentrations of BDNF against H_2_O_2_-induced cell oxidation. On the other hand, in the combination treatment, SH-SY5Y cells were pretreated with H-LIPEF and BDNF for 4 h, and then exposed to 500 μM H_2_O_2_ in the continuous administration of H-LIPEF for another 24 h. In present work, we applied the most effective H-LIPEF parameters (200 Hz, 10 V/cm, and 2 ms pulse duration) previously used by our team for neuroprotection study [Chen et al., 2021] to human neural cells SH-SY5Y. As shown in Fig. 2C, the cell viability decreased to about half of the control group after treating SH-SY5Y cells with 500 μM H_2_O_2_ for 24 h. When cells were treated by combination treatment with H-LIPEF and BDNF, the cell viability was significantly rescued. Under H-LIPEF stimulation, administration of 25 ng/ml and 50 ng/ml BDNF on SH-SY5Y cells increased cell viability to approximately 80% and 90%, respectively. Our results showed that H-LIPEF stimulation demonstrated a positive neuroprotective, while administration of BDNF alone at low doses had a marginally protective effect on SH-SY5Y cells. Noteworthily, the neuroprotective effect was further enhanced by the H-LIPEF and BDNF combination treatment. Our goal is to achieve a better therapeutic effect via H-LIPEF stimulation combined with a lower dosage of drug. Therefore, based on these data, the H-LIPEF (200 Hz, 10 V/cm, and 2 ms pulse duration) parameters in combination with the concentration of 25 ng/ml BDNF were chosen to be used in all the following experiments.

**Fig. 2.**
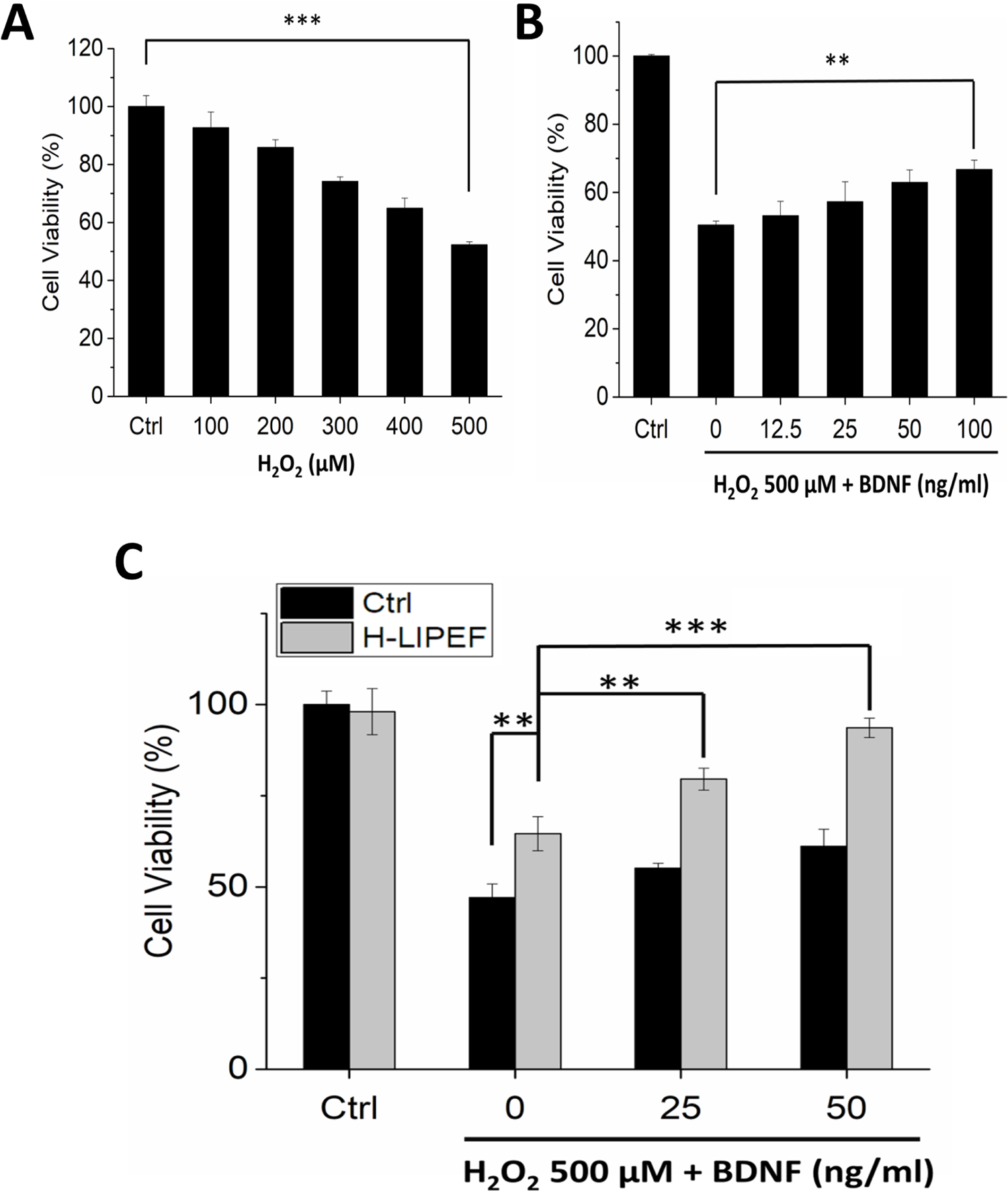
Effect of H-LIPEF and/or BDNF on H_2_O_2_-induced cytotoxicity in SH-SY5Y cells. (A) The concentration response of SH-SY5Y cells treated with different concentrations of H_2_O_2_ and the cell viability was measured by MTT assay at 24 h after the H_2_O_2_ treatment. (B) SH-SY5Y cells were pretreated with different concentrations of BDNFM H_2_O_2_ for 24 h and then the cell viability was measured by MTT assay. (C) SH-SY5Y cells administered by 25 and 50 ng/ml BDNF were subsequently treated with or without H-LIPEF for 4 h and then challenged with 500 μ H_2_O_2_, respectively. The cell viability was also measured by MTT assay at 24 h after the H_2_O_2_ treatment. Data represent the mean ± standard deviation of triplicate tests. The concentration response curve of SH-SY5Y cells was statistically analyzed using one-way ANOVA with Tukey’s post hoc test. **P < 0.01, ***P *<* 0.001.

### Combination of H-LIPEF and BDNF attenuates H_2_O_2_-induced ROS generation

To the best of our knowledge, the damage of ROS accumulation to nerves is known to be the main cause of many brain diseases, such as Alzheimer’s disease and Parkinson’s disease. In view of this, we studied the effect of combination treatment with H-LIPEF and BDNF on the regulation of ROS level by flow cytometry measurements (Fig. 3A). As shown in Fig. 3B, the quantification results show that the ROS level was increased to about 200% of the control level after exposure to 500 μM H_2_O_2_ for 24 h. Compared to the H_2_O_2_ alone group, H-LIPEF treatment significantly ameliorated the H_2_O_2_-induced increase in ROS level, while with BDNF involved, the combination of H-LIPEF and BDNF further enhanced the effect. This result indicates that the combination treatment (H-LIPEF+BDNF) is more effective in reducing the oxidative stress of SH-SY5Y cells.

**Fig. 3.**
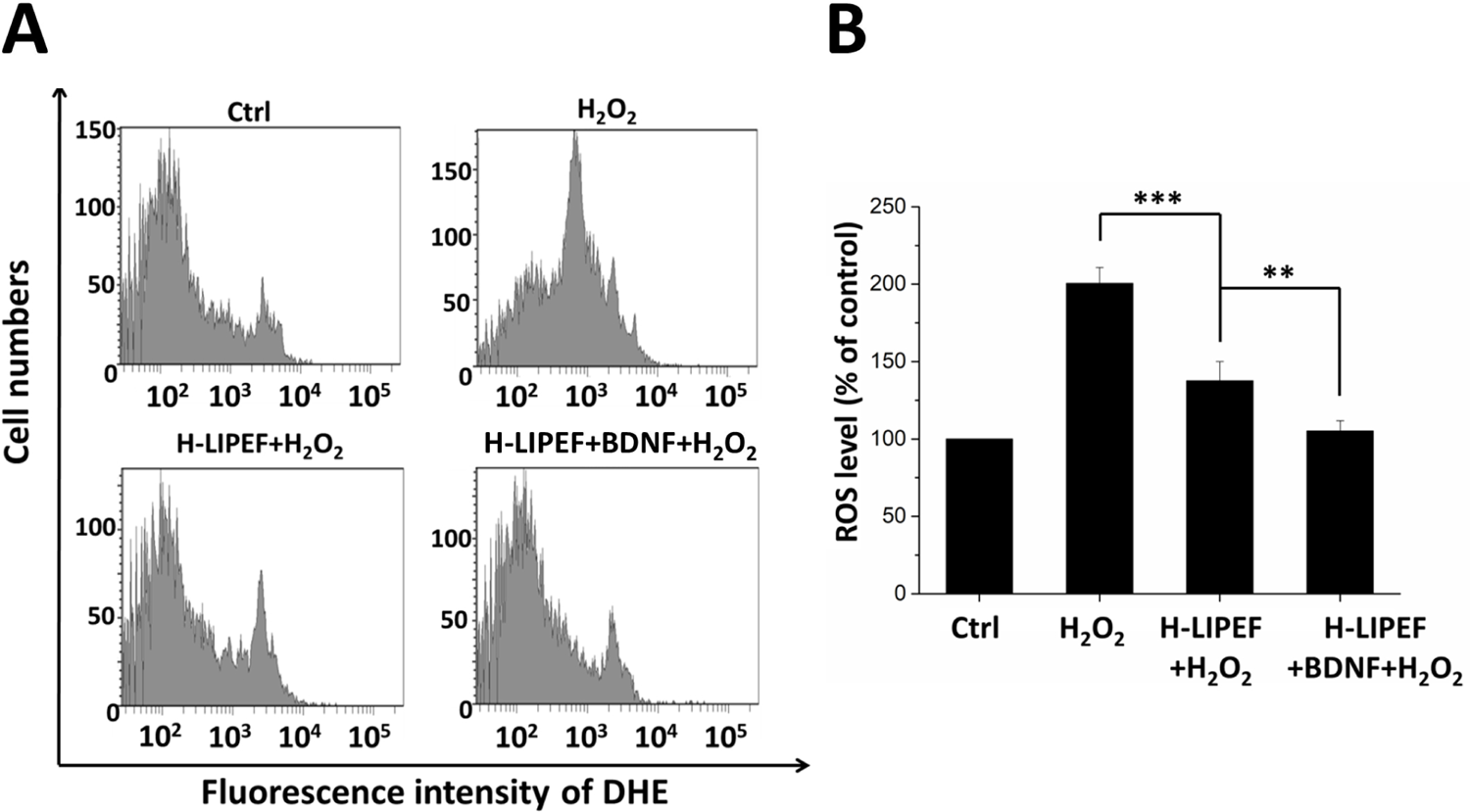
Effect of H-LIPEF alone or in combination with BDNF on H_2_O_2_-induced ROS generation in SH-SY5Y cells. (A) ROS level was measured by flow cytometry with DHE fluorescent dye. (B) Quantification of the ROS levels after H_2_O_2_, H-LIPEF+H_2_O_2,_ and H-LIPEF+BDNF+H_2_O_2_ treatment. Data represent the mean ± standard deviation of triplicate tests. Comparisons of the effect of H_2_O_2_, H-LIPEF+H_2_O_2_, and H-LIPEF+BDNF+H_2_O_2_ on the ROS generation were analyzed using one-way ANOVA with Tukey’s post hoc test. **P < 0.01, ***P *<* 0.001.

### Role of the EGFR pathways in the neuroprotective effect of the H-LIPEF and BDNF

BDNF is an important neurotrophin that regulates the survival of existing neurons and promotes the growth and differentiation of new neurons. It has been previously reported that BDNF can activate EGFR proteins and cause intracellular signalling pathways [Qiu et al., 2006]. Consequently, to further identify whether the protective effect of the H-LIPEF and BDNF on SH-SY5Y cells was related to the EGFR pathway, we examined the protein expression of p-EGFR on human neuroblastoma SH-SY5Y cells. As shown in Fig. 4, when SH-SY5Y cells were treated with H_2_O_2_ alone, the expression of p-EGFR was slightly decreased. In contrast, the H-LIPEF treatment was found to induce a significant recovery of p-EGFR protein level in the cells under H_2_O_2_ oxidative stress. Meanwhile, with the participation of BDNF, the result finds that the combination treatment of BDNF and H-LIPEF can further increase the expression level of p-EGFR protein significantly. The study suggests that H-LIPEF could have an interaction with EGFR, stimulating the initiation of EGFR function, or making EGFR itself more susceptible to phosphorylation. Besides, the binding of BDNF to EGFR would further enhance the neuroprotective effect induced by H-LIPEF on H_2_O_2_-treated SH-SY5Y cells. Subsequently, we studied the downstream signalling pathways that could participate in the neuroprotective mechanism of H-LIPEF and BDNF.

**Fig. 4.**
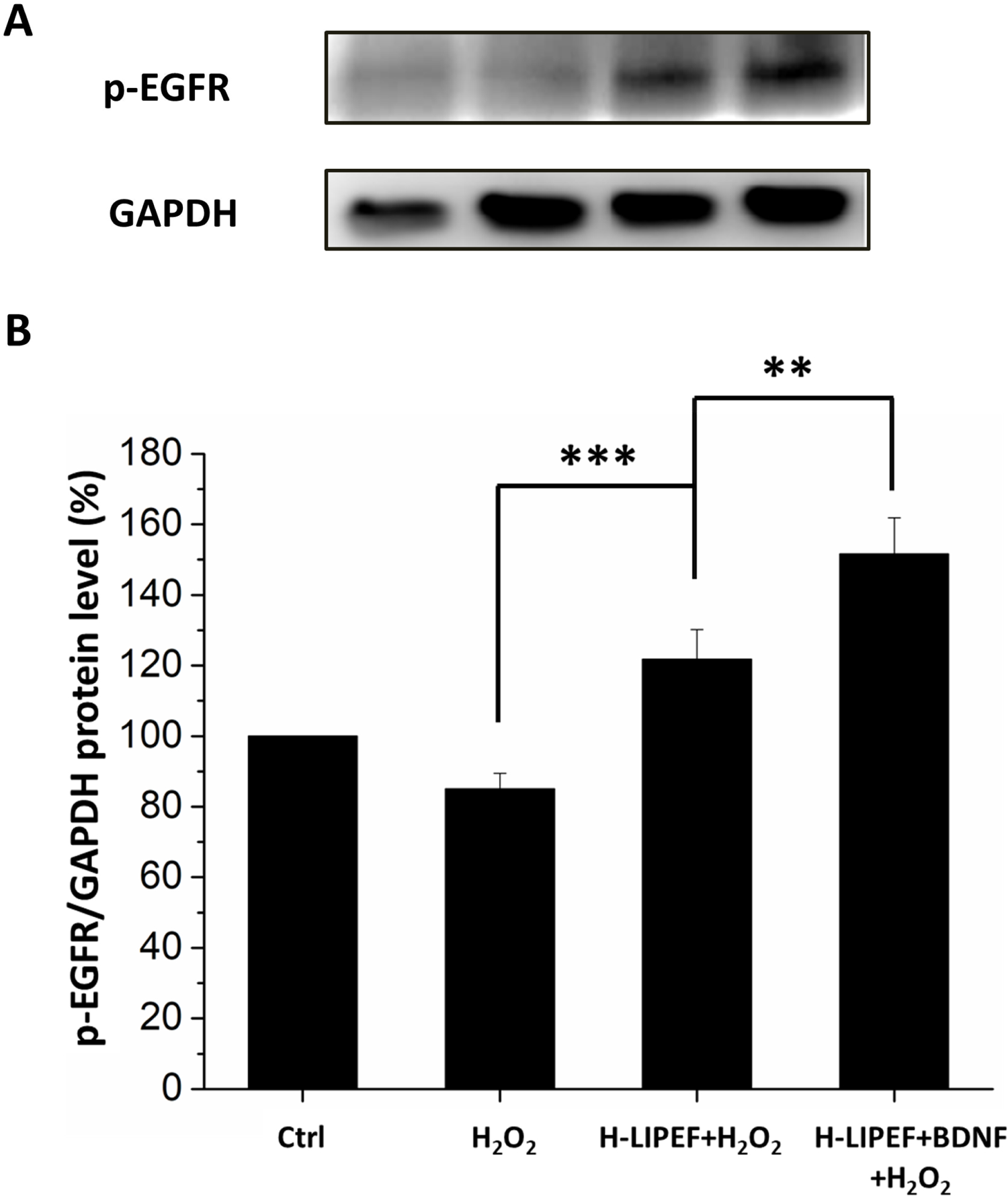
Effect of H-LIPEF alone or in combination with BDNF on expressions of p-EGFR protein in SH-SY5Y cells. (A) Western blot analysis of p-EGFR protein expressions. (B) Quantification of p-EGFR protein expression level after H_2_O_2_, H-LIPEF+H_2_O_2_, and H-LIPEF+BDNF+H_2_O_2_ treatment. H_2_O_2_ and BDNF concentrations in μM and 25 ng/ml, respectively. GAPDH was used as the loading control, and the relative expression level of each group was compared with the untreated control group. Data represent the mean ± standard deviation of triplicate tests. Comparisons of the effect of H_2_O_2_, H-LIPEF+H_2_O_2_, and H-LIPEF+BDNF+H_2_O_2_ on p-EGFR protein levels were analyzed using one-way ANOVA with Tukey’s post hoc test. **P *<* 0.01, ***P *<* 0.001.

### Activation of the ERK pathway in the neuroprotective effect of the H-LIPEF and BDNF

It is well known that EGFR phosphorylation regulates the expression of its downstream proteins such as MAPK family. ERK protein is one of the MAPK family and has been known to be widely associated with cell survival. Upon phosphorylation, activated ERK proteins enter the nucleus, where they can phosphorylate certain transcription factors, thereby regulating the expression of specific genes that contribute to cell survival [Almeida et al., 2005]. As shown in Fig. 5, in comparison to the untreated control cells, the expression level of p-ERK protein was decreased when M H_2_O_2_. Notably, the level of p-ERK was significantly restored in the cells treated with H-LIPEF stimulation. Furthermore, the result found that p-ERK expression was further elevated significantly when SH-SY5Y cells were co-treated by the BDNF and H-LIPEF combination treatment. Therefore, the study indicates that the neuroprotection effect produced by the combination of BDNF and H-LIPEF could be due to the enhanced activation of ERK pathway, thus inducing an even greater recovery of p-ERK level in the cells under H_2_O_2_ oxidative stress.

**Fig. 5.**
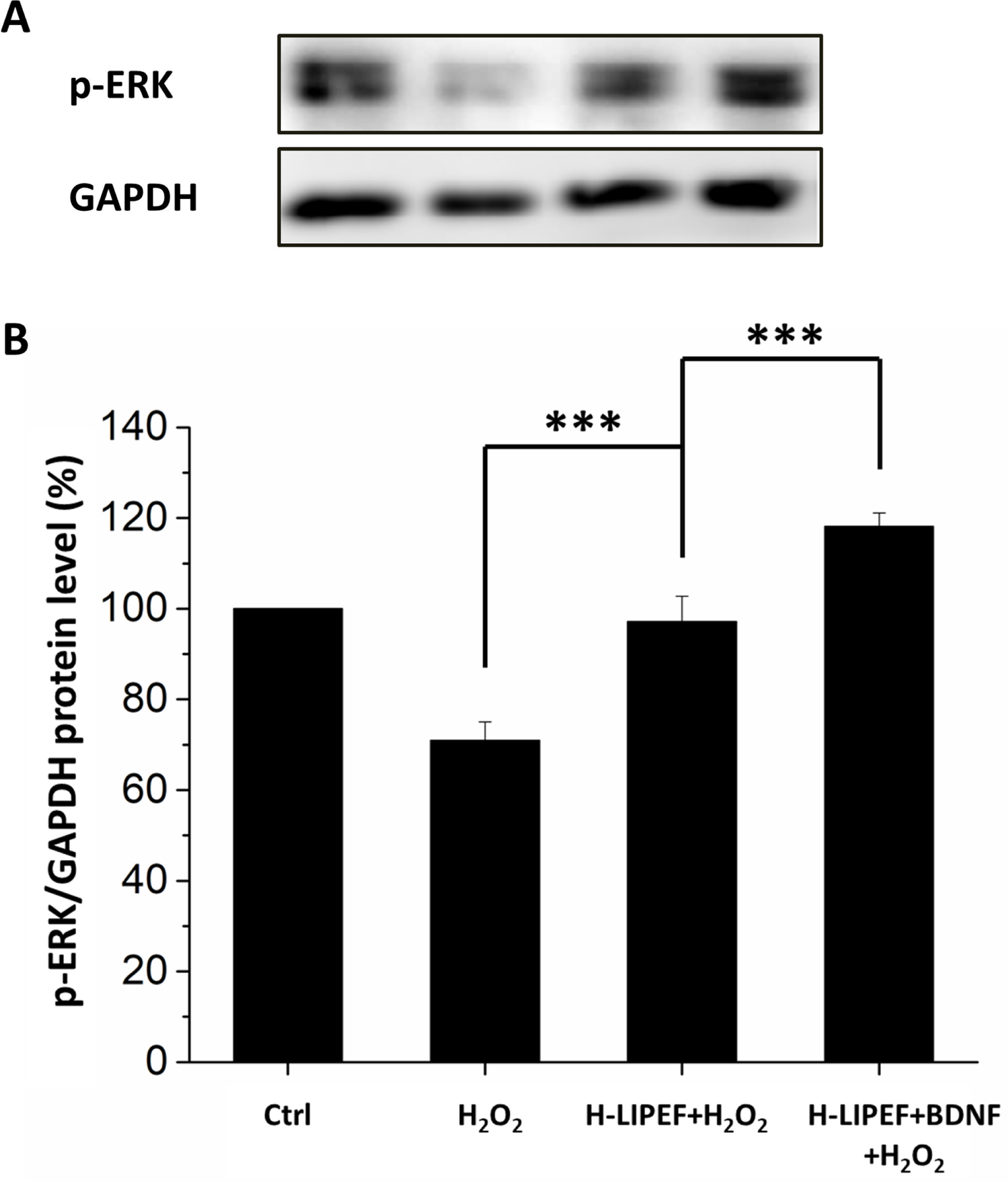
Effect of H-LIPEF alone or in combination with BDNF on expressions of p-ERK protein in SH-SY5Y cells. (A) Western blot analysis of p-ERK protein expressions. (B) Quantification of p-ERK protein expression level after H_2_O_2_, H-LIPEF+H_2_O_2_, and H-LIPEF+BDNF+H_2_O_2_ treatment. H_2_O_2_ and BDNF concentrations in treatments were 500 μM and 25 ng/ml, respectively. GAPDH was used as the loading control, and relative expression level of each group was compared with the untreated control group. Data represent the mean ± standard deviation of triplicate tests. Comparisons of the effect of H_2_O_2_, H-LIPEF+ H_2_O_2_, and H-LIPEF+BDNF+H_2_O_2_ on p-ERK protein levels were analyzed using one-way ANOVA with Tukey’s post hoc test. ***P *<* 0.001.

### Effect of combination of H-LIPEF and BDNF on CREB, Nrf2, Bcl-2, and Bax in H_2_O_2_-treated SH-SY5Y cells

Our previous research pointed out that the protective effect of H-LIPEF on SH-SY5Y cells was through activation of ERK pathway [Chen et al., 2021]. In present study, we examined the effect of the H-LIPEF combined with BDNF on SH-SY5Y cells under H_2_O_2_ oxidative stress. The ERK downstream proteins should be measured to determine whether the combined treatment of H-LIPEF and BDNF can further enhance the ERK signalling pathway. In the following, we further examined the expression levels of p-CREB, Nrf2, Bcl-2, and Bax proteins using western blot analysis. As a transcription factor that regulates the expression of pro-survival proteins, CREB plays an important role in neuronal survival [Sakamoto et al., 2011]. As shown in Fig. 6A, the protein expression of p-CREB in SH-SY5Y cells was decreased when cells were exposed to 500 μM H_2_O_2_. Meanwhile, we found that the expression of p-CREB was restored significantly in SH-SY5Y cells treated with H-LIPEF stimulation. In addition, the combination of BDNF and H-LIPEF was found to further enhance the expression of p-CREB in H_2_O_2_-treated cells significantly. Next, Nrf2 is a key transcription factor that regulates the expression of antioxidant proteins participating in the protection of oxidative damage triggered by injury and inflammation [Ma, 2013]. As shown in Fig. 6B, the level of Nrf2 was also decreased in H_2_O_2_-treated SH-SY5Y cells compared to the untreated cells. In contrast with the cells exposed to 500 μM H_2_O_2_, the expression level of Nrf2 in the cells treated with H-LIPEF was restored significantly. Moreover, the result found that the combined treatment of BDNF and H-LIPEF further increased the expression of Nrf2 in H_2_O_2_-treated cells significantly.

**Fig. 6.**
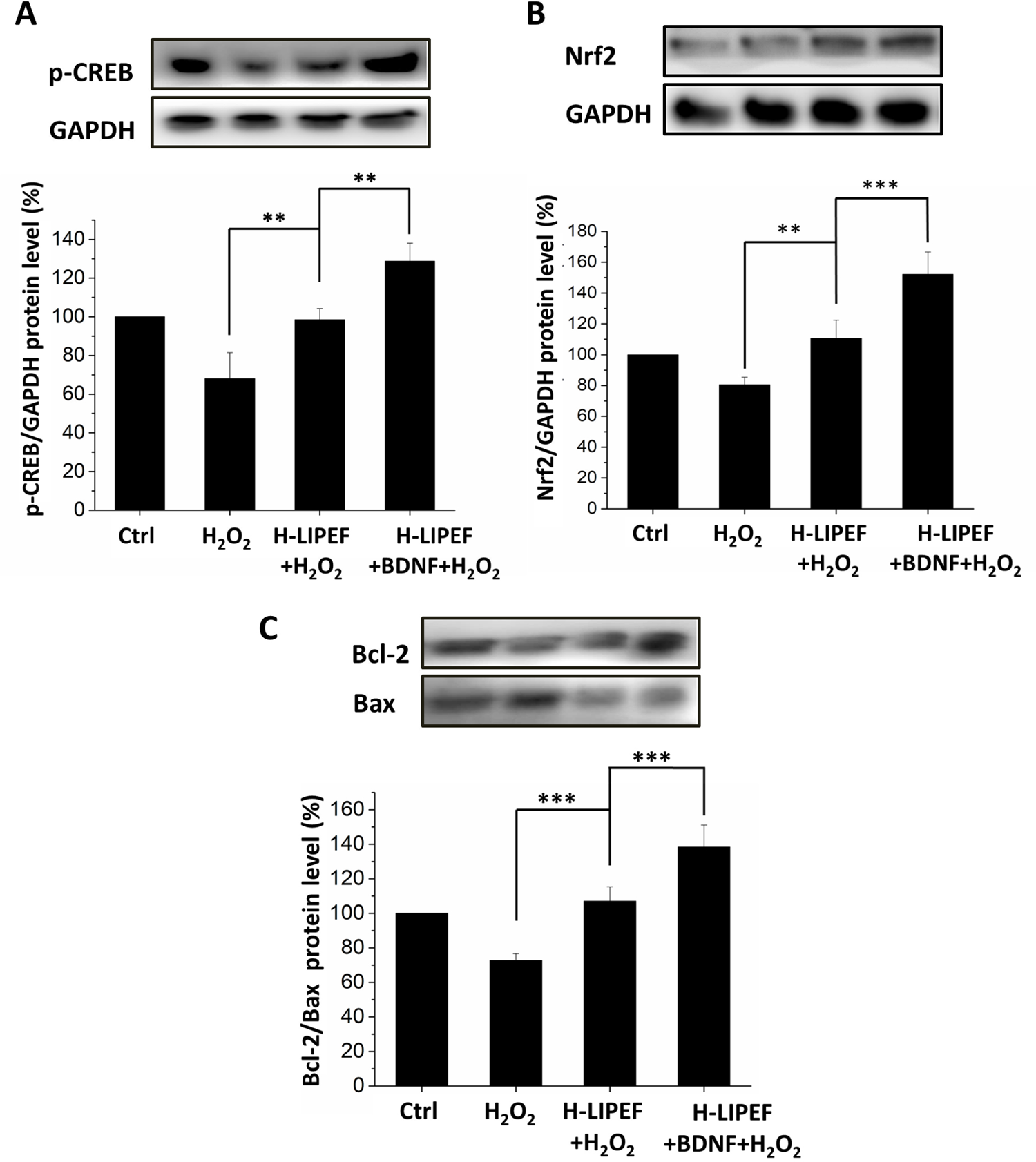
Effect of H-LIPEF alone or in combination with BDNF on survival-related proteins in SH-SY5Y cells. Western blot analysis of p-CREB (A), Nrf2 (B), Bcl-2 and Bax (C) protein expressions. H_2_O_2_ and BDNF concentrations in treatments were 500 μ 25 ng/ml, respectively. The expression levels of p-CREB and Nrf2 were normalized to GAPDH and each relative expression level was compared with control. Data represent the mean ± standard deviation of triplicate tests. Comparisons of the effect of H_2_O_2_, H-LIPEF+H_2_O_2_, and H-LIPEF+BDNF+H_2_O_2_ on different protein levels were analyzed using one-way ANOVA with Tukey’s post hoc test. **P < 0.01, ***P < 0.001.

Furthermore, it is known that both phosphorylated CREB and Nrf2 proteins can regulate the expression of Bcl-2 family proteins [Walton et al., 1999; Niture and Jaiswal, 2012], and the Bcl-2/Bax ratio determines whether the neurons undergo survival or death after being stimulated [Korsmeyer, 1993; Hsieh et al., 2019; Chen et al.,2021]. Therefore, the expressions of Bcl-2 and Bax proteins were also measured in this study. As shown in Fig. 6C, in comparison to the untreated control cells, the level of Bcl-2/Bax ratio of SH-SY5Y cells was found to decrease when cells were exposed M H_2_O_2_. On the other hand, for SH-SY5Y cells treated with the H-LIPEF, the expression levels of Bcl-2 and Bax were increased and decreased respectively, leading to a higher elevation of the Bcl-2/Bax ratio compared to the H_2_O_2_-treated cells. Moreover, it was observed that the combined treatment of BDNF and H-LIPEF further enhanced the ratio of Bcl-2/Bax significantly in the cells under H_2_O_2_ oxidative stress. The above data indicate that the combination treatment not only can greatly increase the expressions of the phosphorylated CREB and Nrf2 proteins, but also can remarkably enhance the ratio of Bcl-2/Bax for better neuroprotection effect, compared to the single H-LIPEF treatment. Consequently, the result suggests that the combination of BDNF and H-LIPEF synergistically promotes the cell survival against H_2_O_2_ oxidative stress via ERK signalling pathway.

## DISCUSSION

The study aimed to investigate the effect of combining BDNF and H-LIPEF in protecting SH-SY5Y neural cells against oxidative stress. In the study, BDNF was combined with the non-invasive and non-contact H-LIPEF with 200 Hz frequency and 10 V/cm intensity, enhancing the survival rate of SH-SY5Y cells under oxidative stress significantly. As one of the most abundant proteins in human brain, BDNF’s importance cannot be overstated. Many studies have pointed out that low-level of BDNF can cause cognitive impairment in neurodegenerative diseases, such as Alzheimer’s disease, which has affected an increasing number of people worldwide [Shimada et al., 2014; Weinstein et al., 2014; Siuda et al., 2017]. Some studies on low endogenous BDNF expression caused by injury or disease have found that employment of exogenous BDNF is effective in protecting neuron cells or brain tissues [Arancibia et al., 2008; Zhang et al., 2015; Xu et al., 2018]. In the study, with H_2_O_2_ as oxidative stress in a cellular model, MTT data show that the combination treatment of H-LIPEF and BDNF significantly increases the viability of SH-SY5Y cells under H_2_O_2_ stress, plus significant improvement in stress-induced apoptosis and suppression of elevated ROS level caused by H_2_O_2_.

As for the neuroprotective mechanism, our previous study showed that the non-invasive H-LIPEF stimulation, within a specific range of electric field intensity and frequency, can effectively increase the survival rate of SH-SY5Y cells under H_2_O_2_ stress to achieve neuroprotection effect [Chen et al., 2021]. The main mechanism is attributed to the ability of electric field to increase the expression of survival-related proteins such as p-ERK and p-CREB, and thus improve the viability of SH-SY5Y cells. In this work, we used H-LIPEF together with BDNF drug to protect SH-SY5Y cells against H_2_O_2_-induced cell injury. This study employed the combination treatment using the electric field with the optimal intensity and frequency identified in our previous study [Chen et al., 2021] and tried to pinpoint more upstream sources of survival signals in SH-SY5Y cells for neuroprotection.

In recent years, people have gradually come to know the importance of EGFR protein in the nervous system, as some studies have found the correlation between neurodegeneration and reduced EGFR expression [Romana and Bucci, 2020]. Increased activation of EGFR may be conducive to neuroprotection. It is known that EGFR has to be bound with a specific ligand such as epidermal growth factor (EGF) for activation and enhancement of the expression of its downstream MAPK family proteins [Wells, 1999]. Moreover, recent literature has found that EGFR can also be triggered by BDNF to activate its downstream signalling pathways [Qiu et al., 2006]. In addition, it has been reported that electric current stimulation alone could trigger a ligand-independent activation of EGFR and even cell migration [ Wolf-Goldberg et al., 2013]. Since activation of EGFR is thought to be associated with cell survival, this study aims to investigate whether the combination of BDNF and H-LIPEF treatment can further enhance the activation of EGFR signalling pathway synergistically to intensify the ability of SH-SY5Y cells to cope with oxidative stress.

While most electrical stimulation applications called for direct contact of electrode plate with tissues and cells, the study employed the non-invasive and non-contact H-LIPEF to stimulate SH-SY5Y cells, demonstrating that H-LIPEF alone can significantly enhance p-EGFR protein expression in SH-SY5Y cells under H_2_O_2_ oxidative stress (Fig. 4). Moreover, a further significant increase in the expression of p-EGFR was observed in the presence of H-LIPEF in combination with BDNF. As shown in Fig. 5, the data also demonstrated that H-LIPEF combined with BDNF significantly boosting the expression of p-ERK, on top of the modulation of other proteins, such as Nrf2 and p-CREB. It is known that Nrf2 is a cellular transcription factor that regulates the expression of various antioxidant proteins. The combination H-LIPEF and BDNF treatment can effectively activate Nrf2 protein to protect neurons via the antioxidant pathways. The other protein, CREB, is also a transcription factor activated by the phosphorylation from ERK, which can trigger an increase in Bcl-2 protein expression. Experimental results also show that the combination of H-LIPEF and BDNF can have a greater effect in raising the Bcl-2/Bax ratio and exerting a pro-survival effect on SH-SY5Y cells.

Recently, the importance of EGFR in neuroprotection has been on the rise, which may be a noteworthy trend for the development of neurodegenerative drugs [Romana and Bucci, 2020]. Besides, the study demonstrates the potential of combining H-LIPEF and BDNF in activating EGFR and related downstream survival proteins, leading to higher survival, which is an approach that may augment drug efficacy and reduce drug dosage. This suggests that the non-invasive H-LIPEF treatment can synergize with BDNF drug to achieve desirable effect at low drug dosage, minimizing side effects. On the other hand, it should be mentioned that conventional electric stimulation was conducted mainly via contact or invasive manner using direct currents, limiting its applications significantly, especially in some organs, such as brain. Therefore, it is believed that the development of non-contact, non-invasive electric field therapy will be inevitable, which can also be further applied in conjunction with drugs for various therapeutic purposes. Previously, our team has achieved synergistic inhibitory effects on cancer cells by combining such electric fields with herbal extracts such as EGCG, CGA, and curcumin, and has also accomplished satisfactory results in repairing mouse motor neuron cell NSC-34, demonstrating the versatility of non-contact, non-invasive electric field stimulations [Hsieh et al., 2017; Hsieh et al., 2018; Lu et al., 2018; Hsieh et al., 2019 Lu et al., 2020]. In application, the electric field’s strength and frequency would vary among different cellular tissues, depending on accompanying drugs, tissue structure, and protein response, which needs further research.

In conclusion, the study demonstrates that H-LIPEF can work together with BDNF, creating a synergy effect in protecting human neuronal cells SH-SY5Y against oxidative stress. The underlying mechanism is believed to involve activation of EGFR and ERK pathway to regulate ROS level, thereby enhancing cell survival. The study also suggests further research on the combination of H-LIPEF and BDNF-mimetic drugs for treating neurodegenerative brain diseases effectively.

## MATERIALS AND METHODS

### Experimental setup for exposure of the cells to non-contact H-LIPEF

The H-LIPEF stimulation device described in our previous study [Chen et al., 2021] was used for exposure of the SY-SY5Y cells to electric field. In brief, the cells were located between two copper flat and parallel electrodes. The pulsed electric signal was produced from a function generator (Agilent 33220A, Agilent Technologies, Santa Clara, CA, USA) and amplified by a voltage amplifier (PZD 700, Trek, Inc., Medina, NY, USA). In the study, consecutive pulses with the electric field 10 V/cm, frequency 200 Hz, and pulse duration 2 ms were applied across the electrodes for the treatment. Cells treated with continuous exposure of non-contact H-LIPEF were kept in a humidified incubator of 5% CO_2_ and 95% air at 37°C.

### Cell culture

The SH-SY5Y cells were purchased from American Type Culture Collection (ATCC, Manassas, VA, USA) and cultured in MEM/F-12 mixture (HyClone; Cytiva, Marlborough, MA, USA) supplemented with 10% fetal bovine serum (FBS) (HyClone; Cytiva), 100 units/ml penicillin (Gibco; Thermo Fisher Scientific, Inc., Waltham, MA, USA), 100 μg/ml streptomycin (Gibco; Thermo Fisher Scientific, Inc.), 1 mM sodium pyruvate (Sigma-Aldrich; Merck KGaA, Darmstadt, Germany), and 0.1 mM non-essential amino acids (Gibco; Thermo Fisher Scientific, Inc.). The cells were cultivated in a humidified incubator containing 5% CO_2_ and 95% air at 37°C, and harvested using a 0.05% trypsin-0.5 mM EDTA solution (Gibco; Thermo Fisher Scientific, Inc.) to prepare for in vitro experiments.

### H-LIPEF and BDNF treatment

BDNF (Alomone Labs, Jerusalem, Israel) was dissolved in distilled water as a stock solution and stored at −20°C. SH-SY5Y cells were pretreated with various concentrations of BDNF and then subjected to H-LIPEF treatment for 4 h. Subsequently, cells were challenged with H_2_O_2_ and put back again to the H-LIPEF stimulation device for further treatment.

### Cell viability assay

The viability of SH-SY5Y cells after indicated treatments was evaluated using the 3-(4,5-dimethylthiazol-2-yl)-2,5-diphenyltetrazolium bromide (MTT) (Sigma-Aldrich; Merck KGaA, Darmstadt, Germany) assay. To perform the assay, the culture medium was replaced with MTT solution (0.5 mg/mL in SH-SY5Y culture medium) and incubated at 37°C for 4 h. The ability of mitochondrial dehydrogenases to reduce the MTT to formazan crystal was used as an indicator of cellular viability. Following MTT reaction, formazan dissolution was performed by using 10% sodium dodecyl sulfate (SDS) in 0.01M HCl. The optical density of each well was measured at 570 nm, with background subtraction at 690 nm using Multiskan GO spectrophotometer (Thermo Scientific, Hudson, NH, USA). The cell viability was determined based on the formazan intensity and expressed as a percentage of the untreated control.

### Western blot analysis

After the treatments, cells were rinsed with phosphate buffered saline (PBS) and were subsequently subjected to lysis on ice for 30 m in lysis buffer (EMD Millipore, Billerica, MA, USA) containing fresh protease and phosphatase inhibitor cocktail (EMD Millipore). The lysates were centrifuged at 23,000 × g for 1 h at 4°C, and the supernatant was obtained for the determination of protein concentration. The protein extracts were loaded in equal amounts into the wells of a 10% SDS-PAGE and transferred to polyvinylidene difluoride (PVDF) membranes (EMD Millipore). The blots were incubated overnight with primary antibodies at 4°C, followed by incubation with horseradish peroxidase-coupled secondary antibodies (Jackson ImmunoResearch Laboratories, Inc., West Grove, PA, USA) at room temperature for 1 h. In this study, the specific primary antibodies against p-EGFR, phosphorylated ERK (p-ERK), nuclear factor erythroid 2-related factor 2 (Nrf2), phosphorylated cAMP response element binding protein (p-CREB), B-cell lymphoma 2 (Bcl-2), Bcl-2-associated X protein (Bax) (Cell Signaling Technology, Inc., Danvers, MA, USA), and GAPDH (GeneTex, Inc., Irvine, CA, USA) were used. All the antibodies were diluted at the optimal concentration according to the manufacturer’s instructions.

Finally, protein bands were visualized with an enhanced chemiluminescence substrate (Advansta, Inc., Menlo Park, CA, USA) and detected with the Amersham Imager 600 imaging system (GE Healthcare Life Sciences, Chicago, IL, USA). The images were analyzed with Image Lab software (Bio-Rad Laboratories, Inc., Hercules, CA, USA). For normalization of proteins, GAPDH was used as an internal control.

### ROS level detection

After the treatments, the SH-SY5Y cells were collected and washed with PBS. The fluorescent dye dihydroethidium (DHE) (Sigma-Aldrich; Merck KGaA) was used to detect the intracellular ROS levels. Cells were resuspended in PBS and incubated with 5 μM DHE dye for 20 min at 37°C in the dark. The fluorescence intensity was measured by FACSCanto II system (BD Biosciences, San Jose, CA, USA) in the PE channel. The ROS levels were expressed as mean fluorescence intensity for comparison.

### Statistical analysis

The results were presented as mean ± standard deviation and performed in triplicate. Differences of statistical significance were determined by a one-way analysis of variance (ANOVA), followed by Tukey’s post-hoc test. P-value < 0.05 was considered to indicate a statistically significant difference. Analyses were carried out using OriginPro 2015 software (OriginLab).

## Acknowledgements

The authors would like to acknowledge the service provided by the Technology Commons, College of Life Science, National Taiwan University for use of flow cytometry system.

## Authorship contributions

Conceptualization: C.Y.C.; Formal analysis: C.Y.C., G.B.L., W.T.C., Y.Y.K., Y.M.C., H. H.L.; Data curation: C.Y.C., G.B.L.; Investigation: C.Y.C., G.B.L., W.T.C.; Writing – original draft: C.Y.C., G.B.L.; Writing – review & editing: C.Y.C.; Supervision: C.Y.C.; Funding acquisition: C.Y.C.

